# Enhancer-gene maps in the human and zebrafish genomes using evolutionary linkage conservation

**DOI:** 10.1101/244475

**Authors:** Yves Clément, Patrick Torbey, Pascale Gilardi-Hebenstreit, Hugues Roest Crollius

**Affiliations:** École Normale Supérieure, PSL Research University, CNRS, Inserm, Institut de Biologie de l’École Normale Supérieure (IBENS), F-75005 Paris, France

**Author notes:** Corresponding authors: YC, HRC.

## Abstract

The spatiotemporal expression of genes is controlled by enhancer sequences that bind transcription factors. Identifying the target genes of enhancers remains difficult because enhancers regulate gene expression over long genomic distances. To address this, we used an evolutionary approach to build two genome-wide maps of enhancer-gene associations in the human and zebrafish genomes. Enhancers were identified using sequence conservation, and linked to their predicted target genes using PEGASUS, a bioinformatics method that relies on evolutionary conservation of synteny. The analysis of these maps revealed that the number of enhancers linked to a gene correlate with its expression breadth. Comparison of both maps identified hundreds of vertebrate ancestral regulatory relationships from which we could determine that enhancer-gene distances scale with genome size despite strong positional conservation. The two maps represent a resource for further studies, including the prioritisation of sequence variants in whole genome sequence of patients affected by genetic diseases.

## Introduction

Enhancers are short DNA sequences that bind transcription factors and contact promoters in cis to activate or repress the transcription of genes into RNA^1^. This control - or regulation - of gene expression by enhancers ensures the fine tuning of mRNA abundance in cells. Disruption of enhancer function has been shown to lead to abnormal gene expression and thus to disease^2-4^. In addition, the majority of variants identified in Genome Wide Association Studies (GWAS) are found outside coding sequences^5^. Together with the observation that many patients remain undiagnosed after genome sequencing because no plausible coding variant can be incriminated^6^, these considerations underscore the importance of identifying enhancers and their target genes to better understand genome function.

Numerous methods have been developed to identify enhancers across entire genomes. Early methods were based on the analysis of the evolutionary conservation of non-coding sequences^7-9^. More recently, the rise of next generation sequencing technologies has enabled large-scale epigenomics projects to map regulatory regions in a genome, e.g. enhancer-associated histone modifications^10,11^, open chromatin regions^12^ or binding of enhancer-associated proteins on the genome^13,14^. Of note, these approaches predict enhancers through indirect evidence for regulatory function, and do not associate predicted enhancers to their target genes. Although choosing the nearest gene is often used as default^15^, the fraction of enhancers regulating their nearest flanking gene is not known. In fact, it is known that enhancers can regulate genes over long distances, sometimes several hundreds of kilobases (kbp) away, sometimes bypassing other genes^16,17^. The classical case of the Shh gene in mouse demonstrates this quite directly as mutations affecting its expression in the intron of the lmbr1 gene located approximately 1 Mb away^16^.

Linking long distance regulatory regions to the genes they regulate is important to study and understand the function of enhancers. Three main categories of experimental methods have been developed to assign enhancers to target genes in a genome-wide manner. The first uses chromosomal conformation capture techniques to identify physical interaction between two loci in the genome^18-21^. The second measures the correlation of transcription activity between non-coding sequences and nearby genes^22^, assuming the two are signatures of a coordinated regulatory function. Finally, non-coding single nucleotide variants (SNVs) can be associated to significant differences in nearby gene expression, thus qualifying as expression QTL (eQTL) that presumably reside within or close to enhancers^23^. These experimental methods are – by definition – specific to the cell-type or tissue where the experiment is carried out and have been applied mostly in human and mouse genomes, while entire group of vertebrate species (e.g. fish) have no available predictions. The use of methods based on evolutionary principles could solve these difficulties, because they do not depend on the specific biological contexts required by experimental assays and are more easily applicable to multiple species.

We previously developed such a method called PEGASUS (Predicting Enhancer Gene ASsociations Using Synteny), a computational method to predict enhancers and their target genes using signals of evolutionary conserved linkage (or synteny)^24^. The rationale underlying PEGASUS postulates that an evolutionary genomic rearrangement would dissociate a cis-acting enhancer from its target gene, and would therefore be deleterious. Negative selection would hence result in the preservation of local synteny between enhancers and their target gene. PEGASUS works in a cell-type or tissue agnostic manner and relies only on the analysis of evolutionary signals. This is of particular interest for regulatory functions important during human development, where experimental assays are limited. This method was originally tested on the human X chromosome followed by experimental validations of more than 1,000 predicted interactions using transgenic assays^24^.

Here, we applied PEGASUS on the entire human and zebrafish genomes to generate two independent genome-wide maps of predicted enhancer-gene interactions. We exploit these resources to uncover evidence for a direct link in the human genome between the number of predicted enhancers associated with a gene and the number of tissues it is expressed in. By comparing these maps, we outline a set of genes with conserved cis-regulation in vertebrates enriched in brain and development functions. We find that the average distance separating predicted enhancer and their target genes scales with genome size, suggestive of weak selective pressure preserving this distance. Finally, our collections of predicted enhancers-gene associations are a valuable resource for the community, represent testable hypotheses that should facilitate genomic studies (e.g. linking transcription factor ChIP-seq peaks to predicted targets) and accelerate the interpretation non-coding variants in whole genome sequences from patient.

## Results

### Enhancers - target genes maps in the human and zebrafish genomes

We predicted enhancers in the human and zebrafish genomes as Conserved Non-coding Elements (CNEs) and applied the PEGASUS method^24^ to predict their most likely target genes. PEGASUS assigns to a CNE the gene(s) within a pre-defined radius (set arbitrarily to 1Mb in both human and zebrafish) with the most conserved synteny (linkage between a gene and its CNE), which we quantify using an evolutionary linkage score (Figure 1a). For the human genome, we first analysed the UCSC 100-way multiple genome alignment restricted to 35 non-teleost fish vertebrates with good genome reconstruction quality (methods) to identify 1,376,482 human CNEs. We applied PEGASUS on these elements and assigned over 95% of these CNEs (1,311,643) to 18,339 human genes (out of 20,342 protein coding genes in the human genome, Figure 1d). Human CNEs cover 2.5% of the genome. For zebrafish, we generated a multiple alignment of 7 teleost fish genomes (methods), leading to the identification of 111,281 CNEs, 50% of which (55,515) could be linked to 17,363 genes (out of 26,427 protein coding genes in the zebrafish genome). These CNEs cover 0.5% of the zebrafish genome (Supplementary Figure 2). The lower sensitivity in identifying zebrafish CNEs can be explained by phylogenetic sampling differences between the two species (see Discussion and Supplementary materials). The majority of CNEs are close to their target genes: the median CNE-TSS distance is 353 kbp in human and 289 kbp in zebrafish (Figure 1b).

**Figure 1:**
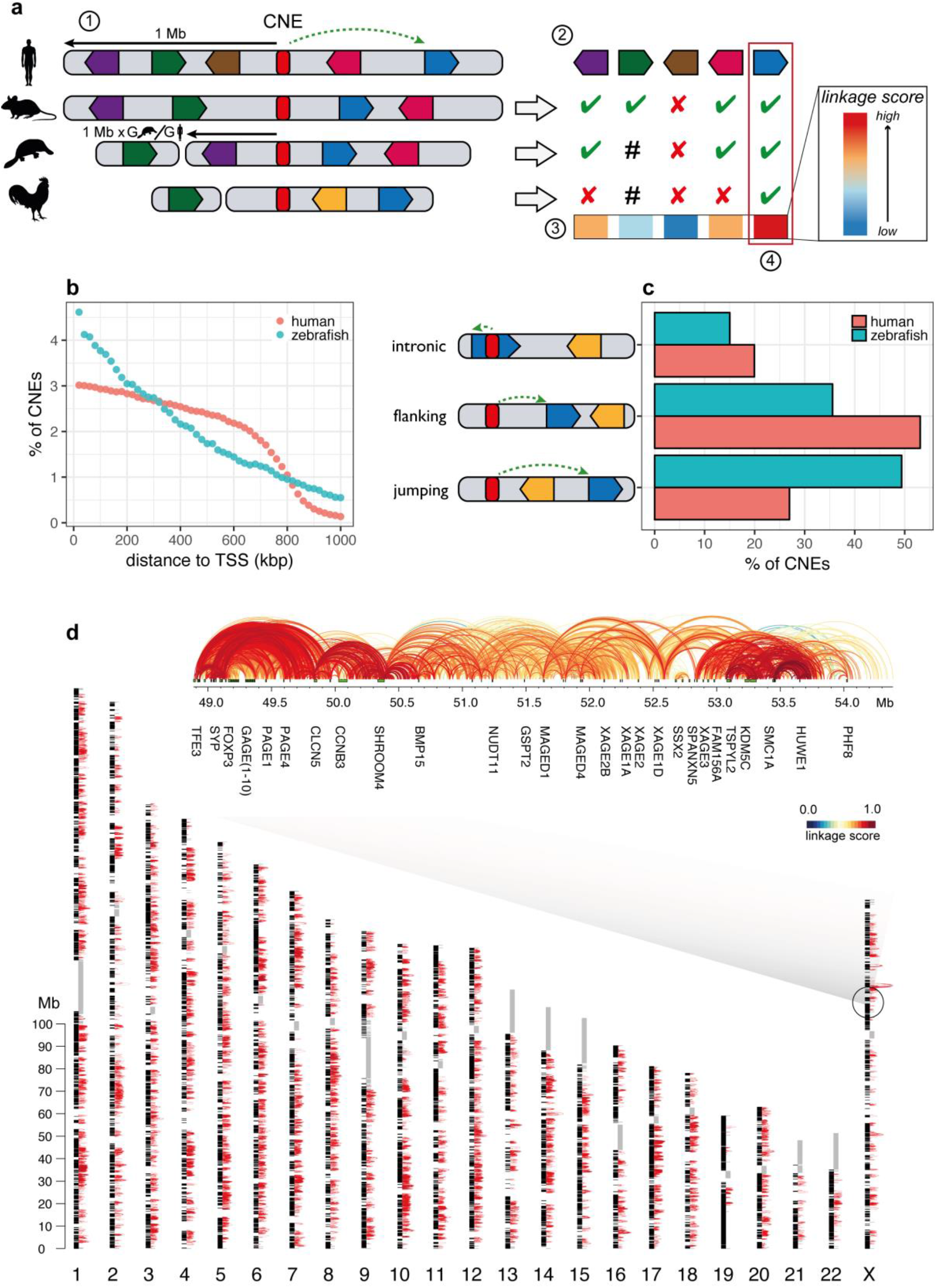
Application of the PEGASUS method and on the complete human and zebrafish genomes. (a) Schematic summary of the PEGASUS method. 1) CNEs (Conserved Non-coding Elements, in red) are identified by cross-species conservation and all genes in a 1 Mb radius are selected as candidate targets. 2) For each gene, the method will look in every species where the CNE was defined if the gene is present in the genome and in the radius (scaled by relative genome size, green ticks), present but outside the radius (hash) or absent from the genome (red crosses). Genes are free to move around within this radius. 3) This information is used to compute a linkage score between a CNE and each gene within a 1 Mb radius. 4) The gene(s) with the highest linkage score is(are) considered to be the most probable target(s). (b) Distribution of CNE-target gene TSS distances. (c) proportion of intronic, flanking and jumping CNEs. (d) Map of CNE-gene interactions in the human genome. For the sake of visibility, only the 174,465 CNE-gene interactions with a PEGASUS score comprised between 0.9 and 1.0 are shown as red arcs. Black blocs alongside chromosomes are protein-coding genes. Grey rectangles are sequences replaced by “Ns” in the hg 19 assembly. An expanded region centred on the FAM71C gene is shown. Green rectangles are protein-coding genes, arcs connect a CNE to the TSS of the predicted target gene and are coloured according to their corresponding linkage score.

The zebrafish enhancer-gene map presented here is the first genome wide resource of its kind. Of note, the human and zebrafish analyses were performed using distinct sets of genomes, enabling rigorous comparisons between phylogenetically independent datasets.

PEGASUS can predict enhancer-gene interactions that skip over unrelated genes. We found that a large fraction of these “jumping” (CNEs skipping over intervening genes to connect to their linked genes) interactions in the human and zebrafish genomes, 27% and 49% respectively (Figure 1c). Moreover, 34% of these “jumping” CNEs in human and 37% in zebrafish are located in an intron of a gene that is not their target gene.

PEGASUS is an *in-silico* method entirely based on evolutionary signals to identify the target genes of CNEs. We evaluated how our predictions coincide with *in-vivo* inferences of regulatory regions (histone modifications^10,25^ or experimental enhancer predictions^22,26,27^). *In-vivo* inferred regulatory regions overlap well our PEGASUS CNEs (up to 95% overlap, up to 4-fold increase over random expectations, Table 1). Finally, we could see a positive link between the PEGASUS linkage score and the overlap between PEGASUS CNEs and in-vivo / in-vitro predicted enhancers & regions with regulatory activity (see Supplementary Material for more details).

**Table 1:** Overlap between in-vitro enhancer predictions and PEGASUS CNEs. *fe*: fold enrichment

| **hg19** |  |  |  | **danRer7** |  |  |  |
| --- | --- | --- | --- | --- | --- | --- | --- |
| source | # of elements | % overlap | *fe* | source | # of elements | % overlap | *fe* |
| Vista (positive)<sup>26</sup> | 846 | 93.6 | 1.7 | differentially methylated regions (all)<sup>27</sup> | 8,225 | 12.9 | 2.6 |
| Vista (negative)<sup>26</sup> | 901 | 91.4 | 1.7 | differentially methylated regions (decreasing)<sup>27</sup> | 1,261 | 30 | 4.6 |
| FANTOM (robust)<sup>22</sup> | 43,011 | 25.7 | 1.9 |  |  |  |  |
| FANTOM (permissive)<sup>22</sup> | 38,554 | 27 | 1.9 |  |  |  |  |
| H3K27ac<sup>10</sup> | 55,728 | 41.4 | 0.9 | H3K27ac<sup>25</sup> | 206,325 | 11.6 | 1.6 |
| H3K4me1<sup>10</sup> | 139,971 | 37.6 | 1.1 | H3K4me1<sup>25</sup> | 226,117 | 11.4 | 1.7 |

We further analysed how PEGASUS *in-silico* target gene assignments coincide with *in-vivo* inferences (conformation capture^19,21^ or expression QTLs^23^). Of note, these *in-vivo* inferences are currently available only for the human genome. We filtered one-to-one overlapping regulatory regions (one and only one PEGASUS CNE overlapping one and only one other regulatory region) predicted to target only one gene and found that between 21% and 27% of target gene assignments were identical between PEGASUS and Capture Hi-C methods and up to 42% between PEGASUS and eQTLs (see Supplementary Material for more details). Interestingly, these numbers were in the same range as what is observed when comparing eQTLs inferences and Capture Hi-C inferences: from 10% to 48% depending on the cell type (10% for ESCs,16% for NECs, 38% for CD34 and 48% for GM12878 cells; see Discussion).

Finally, we show that enhancer-gene associations predicted by PEGASUS are consistent with the 3D organisation of the human genome, because they are located inside topologically associating domains (TADs^28^) more often than expected by chance (proportion tests p-values < 10^−15^, Supplementary Figure 7; see Supplementary Material for more details).

### Genes with more enhancers are expressed in more tissues

Genes cover a broad range of tissue specificities, from ‘housekeeping genes’ required for generic cellular functions and expressed in most tissues to tightly regulated developmental genes sometimes expressed in just a few cells in a short window of time. It naturally follows that the number of enhancers regulating a gene might directly influence the breadth of a its expression pattern, although this has never been demonstrated. We used enhancer-gene interactions predicted by PEGASUS in the human genome to investigate this question, using expression data from the Bgee database^29^. We observe a positive link between the number of adult tissues where a gene is expressed and the number of CNEs targeting this gene (Spearman’s ρ = 0.23, p-value < 10^−15^, Figure 2a).

**Figure 2:**
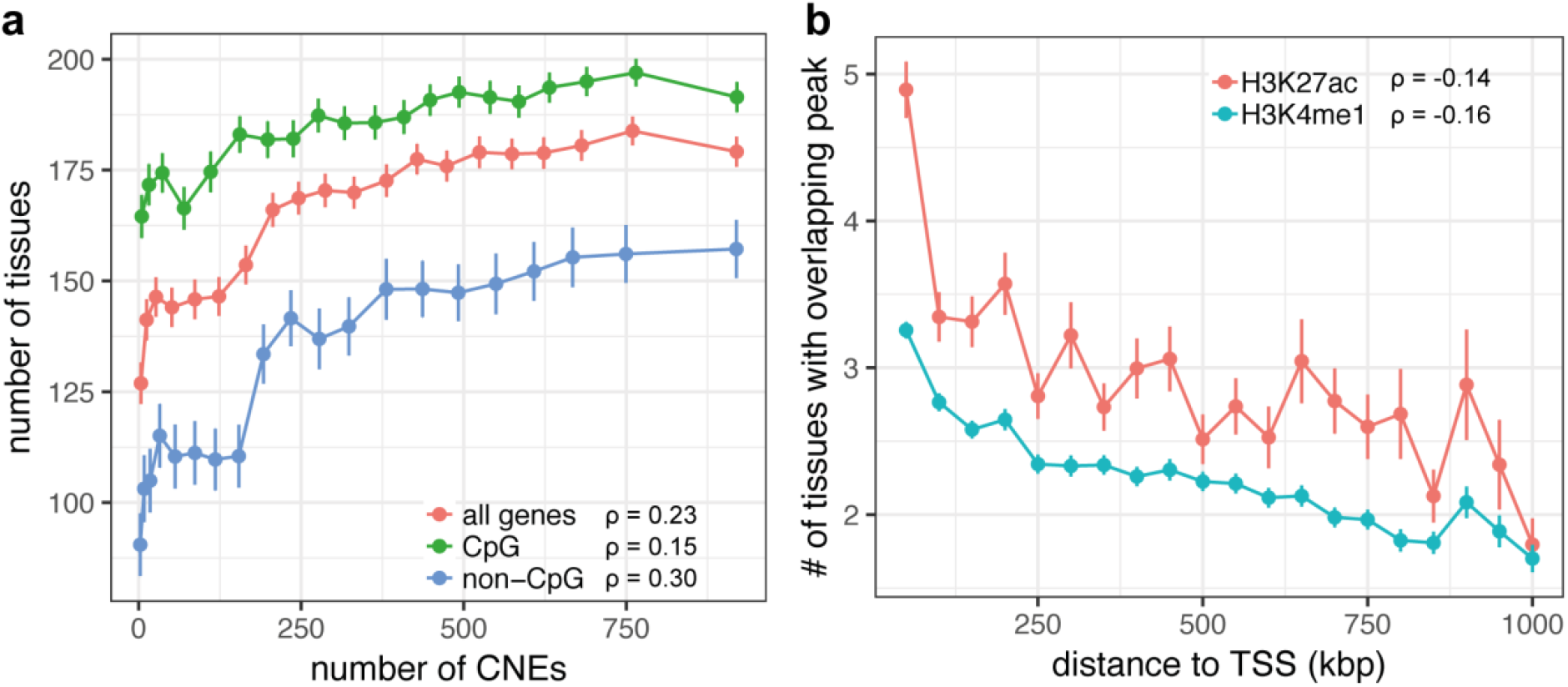
Regulation complexity is positively linked with expression breadth. (a) Link between the number of CNEs targeting a gene and the number of tissues it is expressed in, for all genes or separating between CpG TSS genes and others. Genes were divided in twenty classes of identical size based on their number of CNEs. Classes were made independently for all genes, for CpG genes & for non-CpG genes. Points and vertical lines represent mean number of tissues or life stages with 95% confidence interval in each class. Correlation coefficients were computed on unbinned data. Genes were classified as CpG TSS or non-CpG TSS based on the overlap of their TSS with CpG islands. (b) Link between CNE-TSS distance and predicted CNE activity breadth. CNE activity breadth was computed as the number of tissues for which a CNE overlaps a histone modification ChIP-seq peak. ChIP-seq data comes from the ENCODE project^10^. CNEs were divided in independent classes according to their CNE-TSS distances (one class covers 50kbp). Points and vertical lines represent mean number of tissues with 95% confidence interval in each class. Correlation coefficients were computed on unbinned data.

Genes with promoters falling into CpG islands (regions of elevated CpG content; methods) are usually broadly expressed while other genes are more tissue-specific^30-32^. We therefore sought to further disentangle the link between CNE number and expression breadth by taking the CpG dinucleotides content of gene promoters into account. We split our target genes between those with a transcription start site (TSS) overlapping a CpG island (referred to as CpG genes) and other genes (referred to as non-CpG genes). The correlation between CNE number and number of tissues is positive in both sets of genes. This link appears to be stronger for non-CpG genes than for CpG genes (Spearman’s ρ = 0.30 & 0.15, respectively, both p-values < 10^−15^, Figure 2) but this is explained by the narrower range of tissue where non-CpG island genes are expressed, leading to different ranges of tissue number for both sets of genes (Supplementary Figures 8 & 9).

Conversely to what is observed for genes, CNEs close to their targets tend to be active in more tissues. We computed the correlation between the CNE-TSS distance and the number of tissue a CNE is predicted to be active in, using histone modifications from the Encode project (H3K4me1 and H3K27ac^10^). We found a negative correlation between the number of tissues and the CNE-TSS distance (Spearman’s ρ = −0.16 & −0.14 respectively, both p-values < 10^−15^, Figure 2b).

### Function of regulatory interactions conserved in vertebrates

We defined orthologous CNE-gene associations between the human and zebrafish genomes to study features associated with this conservation of regulatory linkage. Such conserved linkage between enhancers and target genes is consistent with a common origin in the ancestor of Euteleostomi (bony vertebrates), the last common ancestor of human and zebrafish. We identified ∼2,000 CNEs conserved between human and zebrafish (1,986 in human, 1,949 in zebrafish) associated by PEGASUS to ∼600 human-zebrafish orthologous genes (567 human genes, 607 zebrafish genes, see methods). Functional enrichment analyses show that these ancestral regulatory associations are highly enriched in neuronal functions and development (Tables 2 & 3). 30% of these predicted associations are annotated as “jumping” over one or more genes in both species. This includes DMRTA2, a transcription factor involved in female germ cell development^33^ and in brain development^34^, or TSHZ1, a member of the teashirt gene family involved in olfactory bulb development^35^. The strong enrichment in core developmental functions observed with orthologous PEGASUS predictions (Table 3) is consistent with earlier observations, as enhancers identified through sequence conservation are often found to be active during development, especially in the nervous system^7,8,36,37^.

**Table 2:** Top 10 overrepresented anatomy terms (TopAnat^29^) in human genes with conserved regulation with zebrafish. *fe*: fold enrichment. All terms have a false discovery rate lower than 0.002

| **hg19** |  | **danRer7** |  |
| --- | --- | --- | --- |
| *anatomy term* | *fe* | *anatomy term* | *fe* |
| endothelial cell | 1.56 | dorsal thalamus | 8.89 |
| lining cell | 1.56 | blood vessel endothelium | 6.85 |
| barrier cell | 1.56 | cardiovascular system endothelium | 6.49 |
| meso-epithelial cell | 1.56 | pretectal region | 6.47 |
| frontal pole | 1.47 | vestibulocochlear ganglion | 5.98 |
| pole of cerebral hemisphere | 1.47 | preoptic area | 5.94 |
| endothelial cell of viscerocranial mucosa | 1.4 | brain ventricle/choroid plexus | 5.94 |
| buccal mucosa cell | 1.4 | brain ventricle | 5.94 |
| cardiac muscle tissue | 1.39 | ventricular system of brain | 5.94 |
| myocardium of atrium | 1.39 | spinal cord interneuron | 5.92 |

**Table 3:** Top 10 overrepresented Gene Ontology^54^ terms in human genes with conserved regulation with zebrafish. *fe*: fold enrichment. All terms have a false discovery rate lower than 0.05

| **hg19** |  | **danRer7** |  |
| --- | --- | --- | --- |
| *GO term* | *fe* | *GO term* | *fe* |
| ventral spinal cord interneuron differentiation | 14.02 | potassium ion import | 11.28 |
| positive regulation of heart growth | 12.46 | central nervous system neuron differentiation | 7.06 |
| positive regulation of cardiac muscle cell proliferation | 11.89 | embryonic cranial skeleton morphogenesis | 5.88 |
| positive regulation of cardiac muscle tissue growth | 11.6 | cranial skeletal system development | 5.71 |
| central nervous system projection neuron axonogenesis | 11.38 | embryonic skeletal system morphogenesis | 5.71 |
| proximal/distal pattern formation | 9.35 | embryonic skeletal system development | 5.3 |
| positive regulation of organ growth | 9.14 | skeletal system morphogenesis | 5.26 |
| cell fate determination | 8.9 | cell fate commitment | 4.88 |
| positive regulation of cardiac muscle tissue development | 8.63 | skeletal system development | 4.33 |
| regulation of heart growth | 8.06 | positive regulation of transcription from RNA polymerase II promoter | 4.19 |

We validated a predicted ancestral association using a CRISPR-Cas9 mediated knock-out approach. We focused on one CNE of the zebrafish genome, predicted to be associated with a single gene named irx1b. This gene plays multiple roles during pattern formation of vertebrate embryos^38,39^, and we expect its expression pattern to be tightly regulated by a complex array of enhancers. The CNE has evidence for a functional activity during development: it overlaps H3K27ac and H3K4me1 marks as well as ATAC-seq peaks (Figure 3a) and is conserved in all vertebrates. The human orthologous CNE is associated by PEGASUS to IRX1 and IRX2 and also shows evidence for a functional role in this species (H3K4me1 & H4k27ac^10^) as well as sequence conservation. The deletion of the CNE greatly decreases the expression of the endogenous gene in several structures of the zebrafish embryo (Figure 3b,c) establishing it as a *bona fide* developmental enhancer. Interestingly, the CNE targeted by the deletion is closer to another gene, irx4b, without being associated to it by PEGASUS (Figure 3d), yet the expression of this gene is unaffected by the absence of the CNE (Figure 3e). This further illustrates that choosing the nearest gene as a target of a putative enhancer can lead to false predictions and that PEGASUS can distinguish the correct gene target among closely spaced genes.

**Figure 3:**
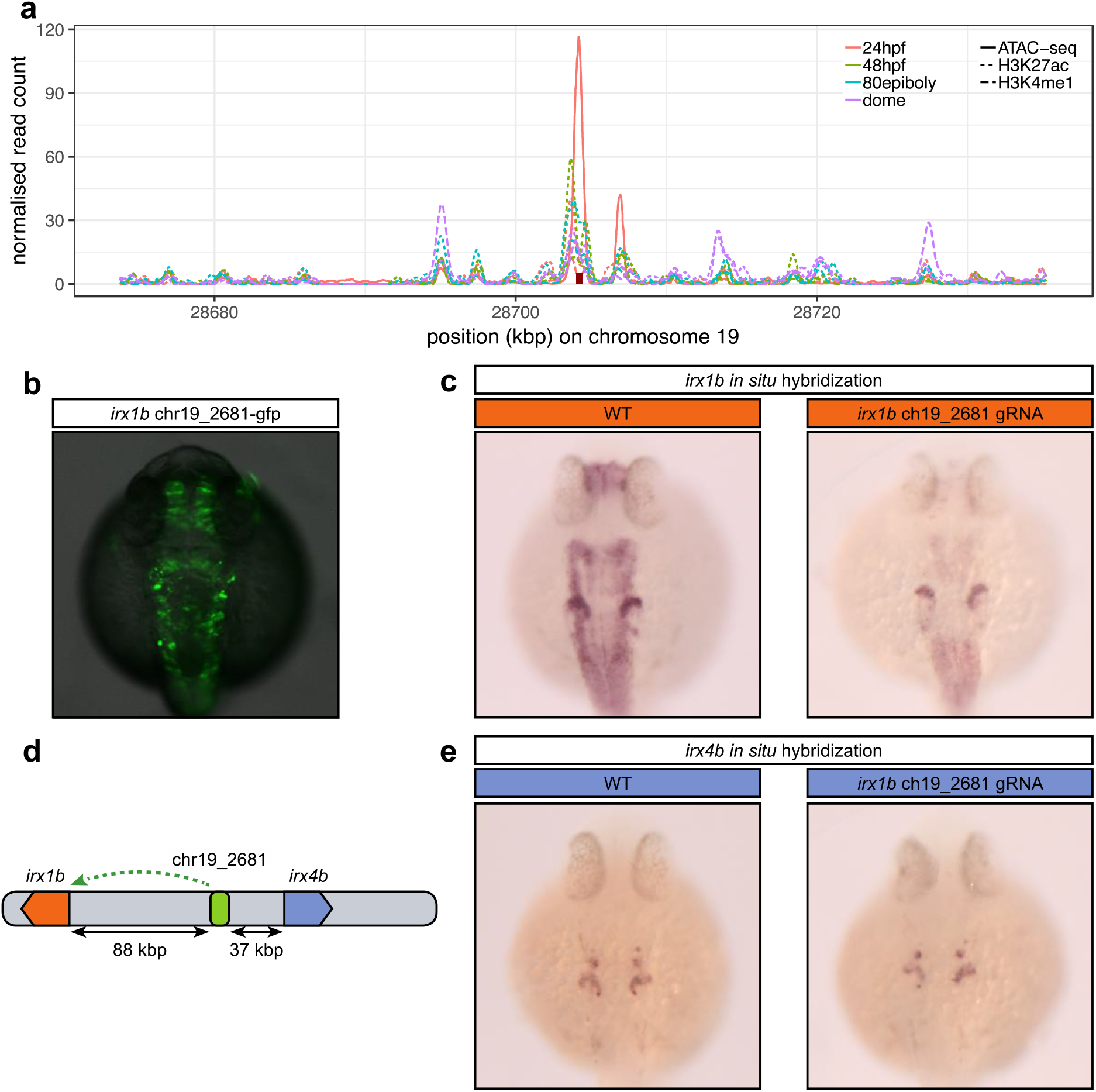
In vivo inactivation of a predicted ancestral enhancer for *irx1b* affects its expression. (a) Evidence for the regulatory potential of the chr19 2681 CNE. The figure shows the normalised read counts for a ChIP-seq analysis of histone modifications (H3K4me1 & H3K27ac) in 4 developmental stages (dashed & dotted lines)^25^ and for an ATAC-seq analysis in 24hpf embryos (full lines)^51^ in a 60kbp region around chr19 2681 (red rectangle). (b) 24h old F0 zebrafish embryos injected with a Tol2 transposon containing the predicted *irx/b* CNE positioned 5’ of the gata2 minimal promoter driving green fluorescent protein (GFP) expression. (c) In situ hybridization for *irx/b* mRNA performed on 24h old wild type embryos (WT) or embryos injected with a mix of 3 CRISPR/Cas9 ribonucleoprotein complexes targeted at the predicted *irx/b* enhancer. The CNE activity profile overlaps with *irx/b’s* expression profile, which comprises the acousticovestibular ganglia, the caudal diencephalon, the tectum, the hindbrain, the spinal cord and the anterior part of the otic vesicle but not in the mid-hindbrain boundary. *irx/b’s* expression level is greatly decreased in all these structures when the CRISPR/Cas9 complex is targeted to the CNE compared to the control, establishing it as a bona fide *irx/b* enhancer. (d) The chr19_2681 CNE, predicted to target irx1b is located closer to irx4b (37 kbp) than to irx1b (88 kbp). (e) In contrast to (b), *irx4b’s* expression profile which includes the anterior part of the otic vesicle and a few cells in the hindbrain is not affected by the CRISPR/Cas9 complex showing that this CNE is specific to *irx/b* and does not regulate *irx4b*.

### Enhancers-gene distances scale with genome size

The “action range” of enhancers is known to encompass a wide span, from within the target gene itself to more than 1 Mb away^16,17^. Importantly, it has been shown that enhancers can change localization within a TAD without affecting downstream gene regulation^40^. Together with results showing high rates of enhancer turnover between species^41,42^, these observations show on specific examples that little selective constraints exist on maintaining enhancers in a specific position relative to their target genes. We test this hypothesis genome wide using the ∼2,000 predicted gene-enhancer associations conserved between human and zebrafish (two genomes with different sizes, 3.1 Gb and 1.5 Gb respectively). We estimated the relative neutral evolution of genomic distances using the sizes of orthologous introns, which are thought to be under negligible size constraint. Results show that distances between orthologous CNEs and their orthologous target genes scale with intron size (median CNE-TSS distance ratio = 2.23, median intron length ratio = 2.39, Figure 4a). There is therefore no evidence for a constraint on CNE-gene distances, which validates our hypothesis. Perhaps surprisingly, despite this absence of selective constraint on interaction distances, we note that the positions of CNEs relative to their target gene TSS (i.e. whether a CNE lies on the 5’ or the 3’ side of the TSS) is highly conserved. We found that more than 91% of orthologous CNEs are located on the same side of their TSS in the human and zebrafish species (30.6% on the 5’ side and 60.9% on the 3’ side, Figure 4b).

**Figure 4:**
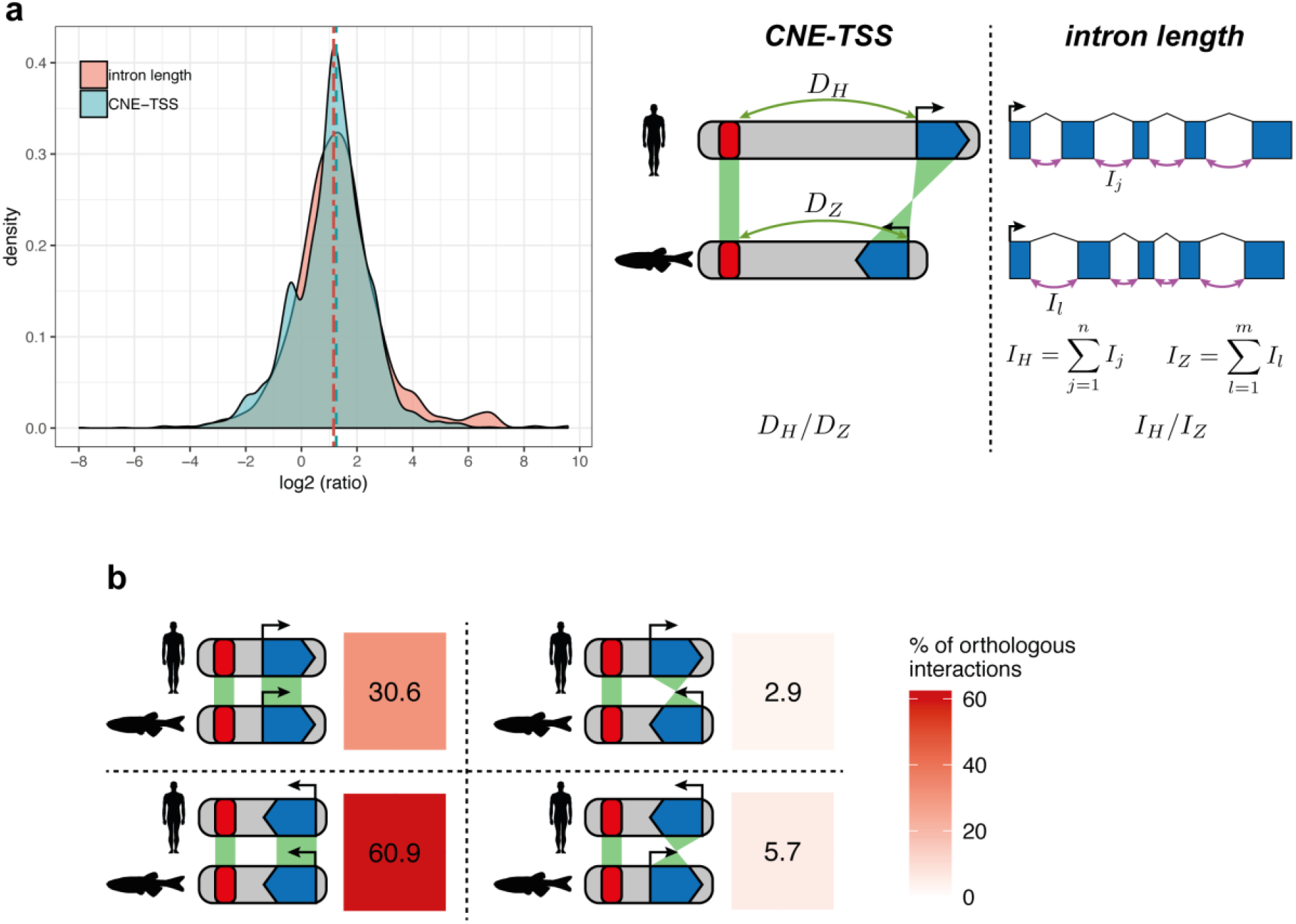
Distances between CNEs and target genes scale with genome size. (a) Pairwise comparison of CNE-TSS distances & intron lengths between human and zebrafish. All comparisons were made using the set of human-zebrafish orthologous genes with conserved CNEs. For enhancer-TSS distances, we compared the CNE-TSS distances (*D_H_* & *D_Z_* for human & zebrafish) for each conserved pair of gene & CNE. For intron lengths, we compared the total intron length (sum of a gene’s intron lengths, *I_H_* & *I_Z_* for human & zebrafish) for each orthologous gene pair. Comparisons were computed as log2(human / zebrafish) ratios. (b) Deep conservation of CNE-TSS relative orientation between human and zebrafish. For the 3570 conserved interactions we analysed, we determined if CNEs were on the 5’ side of the TSS in both species (top left panel), both on the 3’ side of the TSS in both species (bottom left panel), or in different orientations (top & bottom right panels). Numbers represent corresponding percentages of conserved interactions in each category.

## Discussion

We applied the PEGASUS method to identify ∼1,300,000 human and ∼55,000 zebrafish predicted enhancers (conserved non-coding elements) targeting the majority of the genes in their respective genomes. After finding evidence for a regulatory role of these interactions, we show that regulatory interactions ancestral to vertebrates concentrate on core functions necessary to build an organism, that the number of predicted enhancers associated to a gene positively correlates with its breadth of expression and that the distance between predicted enhancers and their target gene evolves neutrally. Our catalogue of enhancer-gene associations contributes to the study of gene regulation by enhancers in vertebrates, can be easily used in a variety of studies and can improve our understanding of gene functions in particular biological contexts.

The first PEGASUS published set of associations was restricted to the human X chromosome^24^. Here, we significantly improve our knowledge of enhancers in vertebrates by applying PEGASUS to the entire human genome, and in the zebrafish genome where no set of enhancer-gene association exists to-date. Moreover, this catalogue can be used to guide and improve the interpretation of epigenomics data such as histone modifications or open chromatin regions or of sequence variants found to be associated to a particular disease in large-scale sequencing projects.

### Effects of phylogenetic sampling

We identified a contrasted number of CNEs between human and zebrafish (∼1,300,000 and ∼55,000, respectively). This discrepancy could be explained by differences in phylogenetic sampling, *i.e*. the number of species and their phylogenetic relationships used for predicting enhancers and linking them to their target gene. Zebrafish was compared to only 6 other genomes, with zebrafish being an outgroup to all but the spotted gar (Supplementary Figure 1 B). In contrast, human was compared to 35 other genomes (Supplementary Figure 1A). We tested the influence of this phylogenetic sampling by comparing the human genome to 6 other genomes that mirror the phylogenetic relationships in the zebrafish study (human being an outgroup to all but one species and equivalent phylogenetic distances as in the fish phylogeny, Supplementary Materials). In this reduced set, we could identify approximately 253,620 CNEs, of which 193,085 (∼82%) target 13,398 genes, a sharp reduction compared to the set identified with a full phylogenetic sampling. The relatively small number of CNEs identified in the zebrafish genome can therefore be explained by the lower number of fish species that can be used for comparative analyses. The addition of more fish species will improve predicted enhancers identifications in the near future.

### The challenges of predicting long-distance regulatory interactions

PEGASUS genome-wide *in-silico* CNE-target gene predictions allow us to directly compare PEGASUS with genome-wide *in-vitro* approaches: we found that PEGASUS predictions and in-vitro predictions agree up to 42% of the time. Most *in-vitro* methods currently employed to predict long-range regulatory interactions in the human genome rely on specific cell lines^19-23^. These methods usually differ in their approach and the cell types or tissues they study, which might limit expectations to observe overlap in their predictions, especially given that many enhancers are tissue-specific^11,20,43^. It is therefore interesting to observe that the level of agreement between eQTLs-based predictions^23^ and Capture Hi-C predictions^19,21^ is equivalent to that observed between these and PEGASUS. In contrast, PEGASUS is agnostic to cell-type or tissue context. The sole rationale underlying the predictions is that the interactions are functional, therefore under sufficient evolutionary conservation to be identified by comparisons with other genomes. Moreover, PEGASUS is able to predict enhancer-gene regulatory interactions that “jump” over one or more genes, which reflects the biology of gene expression regulation more accurately than “nearest gene” approaches. Such “nearest gene” methods are often used to define target genes when studying predicted regulatory regions from epigenomics data (e.g. GREAT^15^). This is a crucial point as experimental methods find a large fraction of these interactions to be “jumping”: between 12% in CD34 cells to 33% in ESCs in Capture Hi-C data, 21% for eQTLs: these interactions will be completely missed by “nearest gene” approaches.

PEGASUS identifies more than 40% of enhancer-gene interactions observed in experimental assays carried out in human cell lines (see Supplementary Figure 4). A much higher overlap may not be expected because the reliance of PEGASUS on evolutionary constraints tend to enrich for interactions active during development^7,13^, and these are typically harder to identify in differentiated cell lines. In addition, given the rapid evolutionary turnover of enhancer regions during evolution^41,42,44^, it is likely that a fraction of cell-type specific enhancers have had little time to leave detectable footprints of selection in a genome. For the same reasons, PEGASUS will fail to capture species-specific or recently evolved regulatory interactions. We observed a general trend for predictions, whether experimental or evolutionary based, to overlap consistently less with increasing distance between the predicted enhancer and the TSS of the target gene (Supplementary Figures 4 & 6) Our data suggests that this could be explained by long distance enhancers being more tissue-specific than short distance enhancers (Figure 2b): short distance enhancers have a regulatory action in more tissues and more likely to have the same predicted target gene when comparing tissues or cell types.

### No evidence for natural selection acting on enhancer-gene distances

Enhancer regulation is mediated through the 3D organisation of the genome. Enhancer-gene interactions occur mostly within TADs^45^, large units of chromosomal interactions largely conserved between cell types and species^28,46^, via DNA looping^47^. Consistent with observations that the distance between an enhancer and its target within a TAD has no effect on its regulatory potential^40^, we show that CNE-TSS interaction evolution between human and zebrafish follows the same pattern as intron size evolution. A recent analysis of genomic regulatory blocks (or GRBs) in metazoans based on the analysis of clusters of conserved non-coding elements showed that these blocks correlate well with known TADs and their sizes seem to correlate well with genome size^48^, providing further evidence that interaction distances between enhancers and target genes are under the same forces that affect genome size in metazoans. Interestingly, our results show that this lack of selective constraint on interaction distances comes with a strong conservation of relative CNE-TSS orientation.

This study provides a unique view of the conservation and evolution of enhancers in vertebrate genomes. Our results based on evolutionary and comparative genomics are complementary to and consistent with genome-wide experimental observations. They support a model where the number of enhancers controlling a gene drives its expression breadth. They also highlight the biological functions with conserved regulation since the vertebrate ancestor. Moreover, the PEGASUS method provides a robust tissue and life stage agnostic target gene prediction method that opens research possibilities in the study of gene regulation in a wide number of species.

## Materials & Methods

### Defining conserved non-coding elements and their most likely target genes

We used a previously published method to predict enhancers and their most likely target genes^24^. This method first predicts enhancers as conserved non-coding elements (or CNEs for short) in multiple genome alignments, and second links a CNE to its most likely target gene(s) as the gene in its vicinity with the most conserved synteny, through the computation of a linkage score measuring this conservation.

We identified CNEs in the human and zebrafish genomes in multiple alignments as follows. We first identified seeds of 10bp with at least 9 alignment columns conserved between all species considered. These seeds were then extended on both sides, allowing up to three non-conserved alignments columns. We allowed up to 40% of mismatches in a column to consider it as conserved for zebrafish and up to 88% for human. For the human genome (GRCH37-*hg*/*9* version), we used the UCSC 100-way multiple alignments restricted to 35 Sarcopterygii species with a scaffold N50 of at least 1 Mb (a full list is available in Supplementary Table 1). Alignment blocks had to contain at least 6 species (including human) with one non-primate species to be considered. For the zebrafish genome *(danRer7/Zv9* version), we generated multiple alignments that include 6 other Neopterygii species (a full list is available in Supplementary Table 2). Multiple alignments were built first by pairwise alignments between zebrafish and other species using LastZ^49^, then by using these to build multiple alignments with Multiz^50^. Alignment parameters can be found in Supplementary Materials. Alignments blocks had to contain at least 3 species (including zebrafish) to be considered.

We used PEGASUS^24^ to identify target genes in both genomes. This method first identifies all protein coding genes (ENSEMBL 75) in a 1 Mb radius around CNEs. It then computes a linkage score for each gene, reflecting the evolutionary conservation of synteny between a CNE and a particular gene. For each gene around a CNE present in *N* species, the linkage score is computed as follows (equation 1 from ^24^):

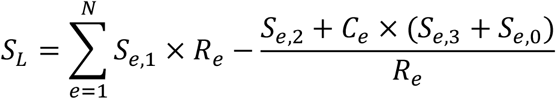

Where *C_e_* is a corrective factor to take assembly errors from low-coverage sequences into account, *R_e_* the rearrangements rate between human or zebrafish and the species *e, S_e,0_, S_e,1_, S_e,2_* and *S_e,3_* the respective status of the orthologous gene considered in species *e* (absent or mis-annotated, present and within the correct radius, present and outside the radius, present and on a different chromosome, respectively). The radius in each species is 1 Mb corrected by the genome size of species *e* normalised by the human or zebrafish genome size. *R_e_* is computed as (equation 2 in ^24^):

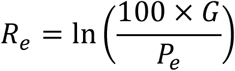

Where *G* is the number of gene pairs in the human or zebrafish genome and *P_e_* the number of these pairs that are direct neighbours in species e. The linkage score is then normalised in a [0,1] interval using a sigmoid transformation (equations 3, 4 & 5 in ^24^). For a given CNE, the gene with the highest linkage score is defined as its most likely target gene. If more than one gene have the highest linkage score, they all are defined as most likely targets. Adjacent CNEs targeting identical gene, present in the same species, having identical linkage scores and distant by less than 100bp were merged together. CNEs located at 100bp or less from an exon were discarded.

### Overlap with functional marks & enhancer predictions

We investigated the link between our PEGASUS predictions and functional marks and previous *in vivo* enhancer annotations. We first computed the overlap with ChIP-seq peaks of histone modifications (namely H3K27ac, H3K4me1, H3K4me3 and H3K27me3) in embryonic stem cells in human^10^ and across various developmental stages in zebrafish^25^. We also computed the overlap between our CNEs and ATAC-seq peaks in zebrafish^51^ and DNase 1 ChIP-seq peaks in human^10^. We finally computed the overlap between our CNEs and other enhancer predictions: we used enhancer predictions from the FANTOM project^22^ or from the Vista database^26^ for human, and from differentially methylated regions^27^ in zebrafish. For all computations, all overlap of at least 1bp were considered. We compared observed overlaps with random overlaps by shuffling CNEs coordinates along the human and zebrafish genomes. Fold enrichments were computed as the ratio between the number of *in-vivo* or *in-vitro*-predicted regulatory regions overlapping PEGASUS CNEs and the number of regions overlapping shuffled PEGASUS CNEs.

### Gene expression data

Gene expression values and calls for the human genome were downloaded from the Bgee database^29^. For each gene, we computed the number of adult tissues for human or developmental stage for zebrafish in which a gene is called as expressed. We filtered out terms (stages or tissues) that had daughter terms for the same gene.

### Comparing target gene predictions with *in vivo* predictions

We compared PEGASUS enhancer-gene predictions in human with other predictions based on eQTLs in various tissues^23^ or on Capture Hi-C data in CD34 and GM12878 cells^19^ and in embryonic stem cell (ESCs) and ESC-derived neurodectodermal cells (NECs)^21^. The latter contains interactions present in one cell type or in both simultaneously (labelled “Both”). We considered only CNEs with a one to one overlap (one and only one PEGASUS CNE overlapping one and only one other predicted enhancer), and both targeting only one gene. In this set of PEGASUS enhancers and other enhancers, we computed for each cell type the percentage of PEGASUS predictions and other prediction that agreed with each other.

### Defining orthologous enhancers & target genes between human & zebrafish

We downloaded human-zebrafish and zebrafish-human pairwise chain alignments from UCSC. We defined orthologous CNEs as human and zebrafish CNEs that overlapped by at least 10bp on either pairwise alignment. We next downloaded human-zebrafish orthologous genes from the Ensembl database (version 75)^52^ to identify orthologous enhancers targeting orthologous genes.

Because of the evolutionary distance between human and zebrafish, some orthologous regions are difficult to align and are thus impossible to detect. To circumvent this problem, we used the spotted gar genome^53^ to identify additional orthologous CNEs. We downloaded human-spotted gar pairwise chain alignments and used our custom-made zebrafish-spotted gar pairwise chain alignments to respectively map human and zebrafish CNEs onto the spotted gar genome. We considered human and zebrafish CNEs as orthologous if they overlapped by at least 10bp on the spotted gar genome. No information other than orthology of CNEs on the spotted gar genome was used. We identified orthologous targets by looking at the orthologous genes set used above. Orthologous CNEs identified both directly and via the spotted gar were combined.

### Gene enrichment analysis

We performed anatomical terms enrichment analyses using the TopAnat webtool of the Bgee database^29^ and the PantherDB webtool^54^. The test set was defined as human-zebrafish orthologous genes with conserved CNEs defined above. The control set was defined in both species as all genes targeted by at least one CNE.

### Distance to transcription start sites

We downloaded transcription start sites (or TSS) locations from the Ensembl database (version 75)^52^. For each gene, we considered only the transcript giving the longest protein. We computed for each enhancer-gene the distance to the TSS as the shortest distance from enhancer boundary to the target’s TSS.

### CNE activity breadth prediction

We computed each CNE’s activity breadth by computing in how many tissues a particular CNE overlaps a histone modification ChIP-seq peak. We focused on H3K27ac and H3K4me1, using ChIP-seq data from the ENCODE project^10^.

### Topologically associating domains

We downloaded topologically associating domains (or TADs) coordinates for two cell types, human embryonic stem cells (hESCs) and IMR90 fibroblasts^28^. We converted these coordinates from *hg/8* to *hg/9* using the liftOver utility available at the UCSC genome browser^55^. For CNEs targeting only one gene, we computed for each cell type whether both an enhancer and its target gene were located within the same TAD. We also computed random overlap between TADs and regulatory interactions by shuffling the localisation of the TAD domains along the human genome and computing the overlap between PEGASUS interactions and these shuffled TADs.

### In vivo validation

#### Vector and cloning

The predicted Irx1b CNE (chr19 2681: chr19:28704114-28704349, *danRer7* version of the zebrafish genome) was amplified from zebrafish genomic DNA using the following primers: CNE-Irx1b-Forward: 5’-TGAATGCTCATCCGGAACATCCACTGCTGCTCCCAAAG-3’; CNE-Irx1b-Reverse: 5’-GACCTGCAGACTGGCAGTTCCTCGCCAGAGCTCAG-3’ and cloned into pZED plasmid^56^ upstream of the minimal GATA2 promoter/GFP reporter.

#### Zebrafish egg injections for transgenesis

The Tol2 transposon/transposase method of transgenesis^57^ was used with minor modifications. Two nanoliters containing 20 ng/μl of transposase mRNA and 30 ng/μl of phenol/chloroform purified pZED construct were injected in one-cell stage embryos.

#### In situ hybridization

In situ hybridization were performed as described^58^, using an *Irx/b* probe corresponding to exon 2.

#### Zebrafish egg injections for mutagenesis

Three RNAs targeting three ultra-conserved sequences in the CNE were designed as follows: CNE-Irx1b-guide1: TCCGTCACGCTGAGATAATC;CNE-Irx1b-guide2: TCAAACACTTTGGGGAACAA;CNE-Irx1b-guide3: TGACCTCTCACCTCGGGCTA. Similarly, three RNAs targeting three ultra-conserved sequences in a random genomic region were designed as follows: Control-guide1: TTGCTTCTGCGCTGAAATAA; Control-guide2: ATGGACTAAAAATTTCACTT; Control-guide3: GAATGTTGATTGTAATTACA. They were purchased from Integrated DNA Technologies as “crRNA”, hybridized with their “tracrRNA”, forming the guide RNA (gRNA) and incubated with a Cas9 protein (gift from J - P. Concordet). Three nanoliters containing a mix of the 3 resulting ribonucleoproteins (Cas9/gRNA) targeting either the control or the predicted *Irx/b* enhancer were injected at l5μM each.

34 embryos showed a signal for decreased gene activity over 37 embryos tested.

## Supporting information

## Data availability

PEGASUS predictions for the human genome *(hg/9)*, the zebrafish genome *(danRer7)* as well as interactions predicted to be conserved between both genomes are available here: ftp://ftp.biologie.ens.fr/pub/dyogen/PEGASUS/

## Acknowledgements

We thank Pierre Vincens for the coordination of computing resources, Morgane Thomas-Chollier and Camille Berthelot for fruitful comments and remarks on earlier versions of this manuscript. This work was supported by grants from the French Government and implemented by ANR (ANR-14-CE13-0004-02, ANR-10-BINF-01-03 Ancestrome, ANR-10-LABX-54 MEMOLIFE and ANR-10-IDEX-0001-02 PSL* Research University).

