## Supplementary material for "Enhancer-gene maps in the human and zebrafish genomes using evolutionary linkage conservation"

### Enhancer-gene maps in the human and zebrafish genomes unravel ancestral vertebrate regulation and principles of enhancer function – Supplementary Information

#### Supplementary Methods: alignment parameters

Zebrafish-centred multiple alignments were build using LastZ [1] and Multiz [2]. Genomes used are indicated in Supplementary Table 2.

We aligned zebrafish to all six species using the same parameters. Alignments parameters were as follows: step=1, masking=0, seed=12of19 (with transitions allowed), hsp threshold=3000, ydrop=3400, gapped threshold=3000, inner=0, with gap opening and extending gaps of 400 and 30. The standard HoxD55 score matrix was used for scores. The chaining step was performed on alignments with a minimum score of 0 with a loose linear gap matrix. Chained alignments were processed into nets from which best chain alignments were extracted (as indicated on the UCSC website).

Pairwise alignments were merged together using Multiz. All blocks (all parameter) of minimum length 1 (R parameter) were kept.

#### Supplementary Materials

##### Effects of phylogenetic sampling on CNE identification.

Predictions gave contrasted number of CNEs between human and zebrafish (~1,300,000 and ~111,000, respectively). Contrasted phylogenetic sampling, i.e. the number of species and their phylogenetic relationships used for CNE identification and target gene predictions, between the two species may cause this discrepancy: for zebrafish, we compared it to only 6 other genomes, with zebrafish being an outgroup to all but the spotted gar (Supplementary Figure 1B). In contrast, for human we compared it to 35 other genomes spread across the phylogeny (Supplementary Figure 1A). We tested the effects of species number and relationship by generating a “control” set of human CNEs and target genes, by comparing human to 6 species that mirror the phylogenetic relationships in zebrafish. The species used were as follows: zebra finch, mallard duck, chicken, turkey, anole lizard and xenopus (see Supplementary Table 1). We run PEGASUS with the same parameters as the original zebrafish set.

In this control set, we could identify 235,620 CNEs, of which 193,085 (82%) could be associated to a target gene, a sharp reduction compared to the main set. We still identify more CNEs in this control set compared to the zebrafish analysis, which could be due to the smaller genome size of fish in general (e.g. 350 Mb for tetraodon, 1.5 Gb for zebrafish, Supplementary Table 2).

#### CNEs and PEGASUS assignments overlap enhancer predictions and experimentally defined regulatory interactions

Predicted enhancers are solely defined by sequence conservation, yet all predicted enhancers are assigned a linkage score as long as a gene lies within a 1 Mb radius. However, not all predicted enhancers are regulatory enhancers and we wished to confirm that the linkage score is correlated with evidence for a regulatory role in the human and zebrafish genomes, as noted previously for human chromosome X [3]. CNE linkage scores are strongly correlated in both species with chromatin marks (H3K27ac, H3K27me3, H3K4me1 and H3K4me3) in human embryonic stem cells [4] or in various developmental stages in zebrafish [5] (Supplementary Figures 3). Interestingly, the two marks associated with active enhancer function (H3K4me1 and K3K27ac [6]) show the strongest overlap. A strong correlation between CNE linkage scores and open chromatin marks (ATAC-seq in zebrafish and DNase1 marks in human) further supports the regulatory role of linked CNEs (Supplementary Figure 3). In addition, CNEs for which no target gene could be identified have almost no experimental evidence for a regulatory function, as expected if PEGASUS correctly classifies those CNEs has playing no role in regulating the expression of a target gene (Supplementary Figure 3). Finally, the overlap between CNEs and experimentally verified enhancers in human [7,8] and zebrafish [9] follows the same notable enrichment profile as a function of the linkage score (Supplementary Figure 3).

We sought to measure the agreement between PEGASUS target gene predictions and those from tissue or cell-type dependent experimental approaches. We restricted our analysis to human CNEs linked to a single gene and with a one-to-one overlap with enhancer predictions with other methods (i.e. one and only one CNE overlapping one and only one enhancer region). When confronted with Capture Hi-C based predictions [10,11], the level of agreement depends on the cell type: from 21% for (NECs) to 27% for GM12878 (a lymphoblastic cell line). The level of agreement with eQTLs-based predictions from the GTEx consortium [12], is higher, with a 42% overlap. Interestingly, for all target gene predictions methods that we analysed, the agreement decreased with distance to the TSS (Supplementary Figure 4), even in the range 0-500 kb where the PEGASUS linkage score is relatively constant (Supplementary Figure 5). This could be explained by the observation that enhancers predicted at longer distances are active in fewer tissues (Supplementary Figure 6), limiting the possibility of overlap with datasets obtained on a limited number of cell or tissue types. The level of agreement between PEGASUS predictions and methods based on functional assays is within the same range as when the latter are compared to each other (between 8 and 38% of eQTLs-based predictions agreed with Capture Hi-C predictions, depending on the cell type), and here also, the overlap decreases with the distance to the TSS (Supplementary Figure 6). We therefore conclude that PEGASUS enhancer-gene predictions are consistent with results from genome wide functional assays. These results are consistent with the fact that PEGASUS, which works in a cell-type or tissue agnostic way, is as powerful in predicting target genes of putative enhancers as labour-intensive methods such as Capture Hi-C. Moreover, using CNEs as putative enhancers enables us to detect more regulatory regions and interactions in a single analysis than experiment-based methods.

#### Long-range regulatory interactions within TADs

Topologically Associating Domains (TADs [13]) have been shown to coincide well with the regulatory landscape governing gene expression [14,15]. Here, for CNEs linked to a single gene, 57% and 66% of predicted interactions indeed reside within a TAD in hESCs and IMR90 cells respectively [13], compared to an average of 32% and 41% respectively when we shuffle TAD intervals (proportion test p-values < $10^{-15}$ for both cell types). CNEs are linked to their target gene with a higher score when inside the same TAD (mean scores, 0.72 inside vs 0.67 outside for hESCs, 0.71 vs 0.68 for IMR90, Wilcoxon rank sum test p-values < $10^{-15}$ for both cell types, Supplementary Figure 7), overlapped more with functional marks (14% vs 10% for H3K4me1 & 15% vs 11% for H3K27ac in hESCs, 13% vs 10% for H3K4me1 & 10% vs 15% for H3K27ac in IMR90, all proportion tests p-values < $10^{-15}$) and were also closer to each other (median distance to TSS, 332 kb inside vs 522 kb outside for hESCs, 355 kbp vs 524 kbp for IMR90, Wilcoxon rank sum test p-values < $10^{-15}$ for both cell types, Supplementary Figure 7). Finally, we see a striking link between the across species conservation of CNEs and their localisation within a TAD. First, human-zebrafish orthologous CNE-target gene pairs (human-zebrafish orthologous genes with conserved CNEs) are more often located within a TAD than expected by chance (proportion tests p-values < $10^{-15}$ for both hESCs and IMR90 cells). More importantly, we see a positive link between the conservation depth of CNEs and the co-localisation of CNEs and target genes within the same TAD (Supplementary Figure 7). The association between TAD co-localisation and both distance to TSS and conservation depth cannot be explained by chance alone (Supplementary Figure 7). These results, in line with previous observations [16], show evolutionary conservation of linkage between CNEs and their target genes is consistent with topological organisation of chromatin.

#### Supplementary Figures

A

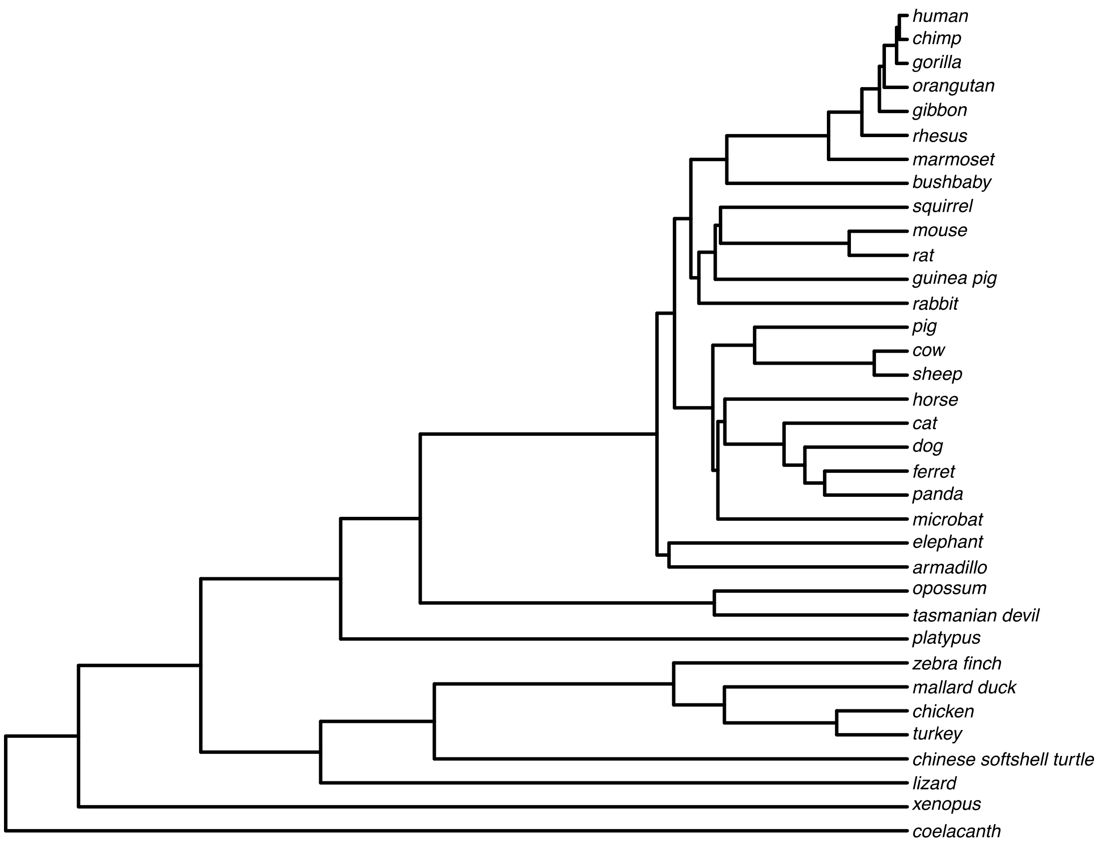


B

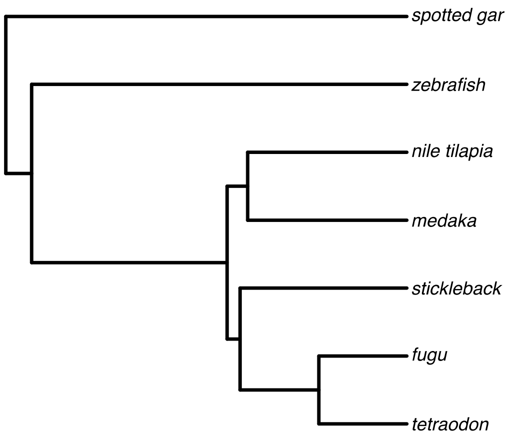


**Supplementary Figure 1:** (A) Phylogenetic relationship for the species used for the human, adapted from the tree used by the UCSC genome portal [17]. (B) The zebrafish tree was computed on a random set of 50 1-to-1 orthologous proteins using PhyML [18]. Both trees were made ultrametric.

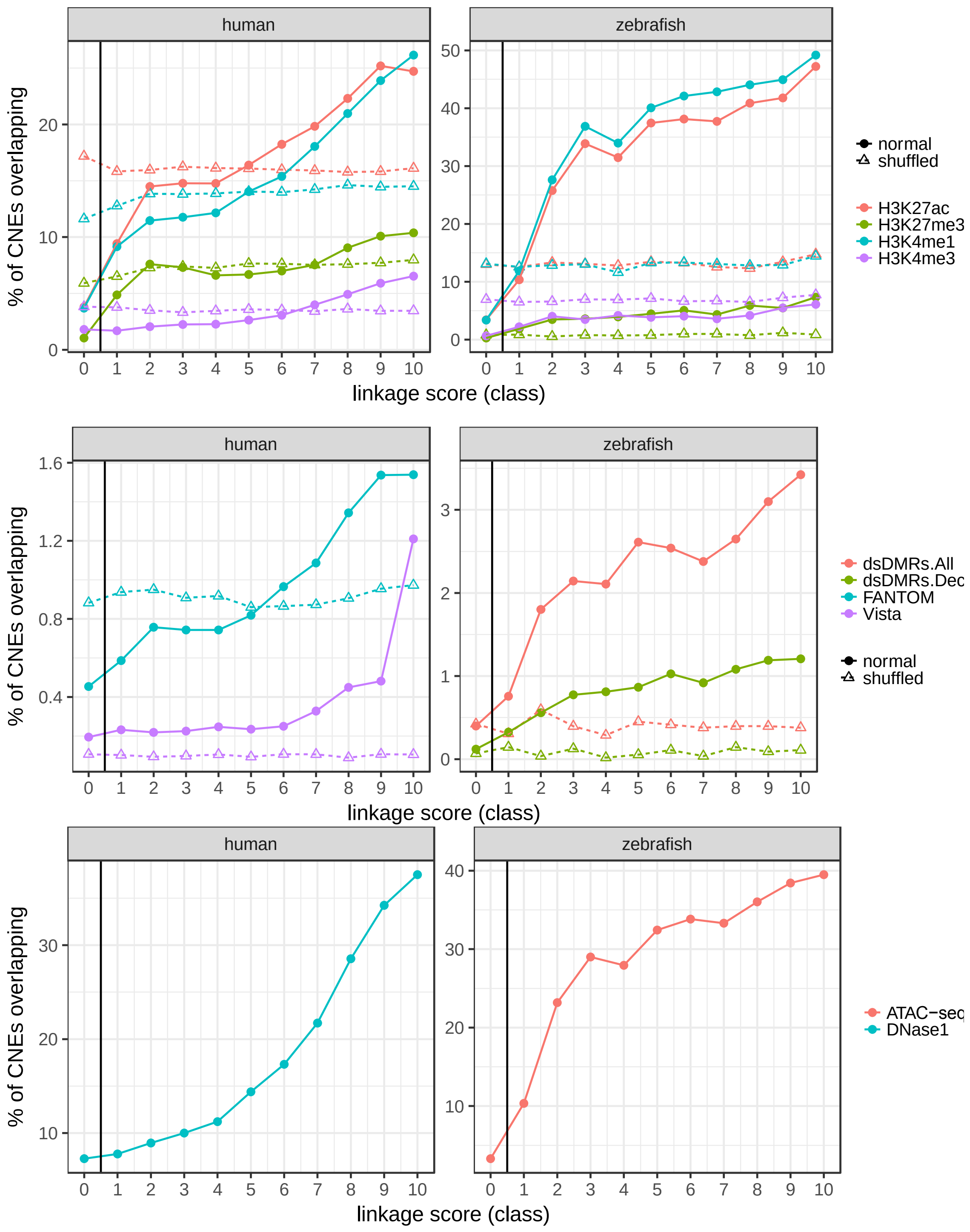


**Supplementary Figure 3:** Percentage of predicted enhancers overlapping histone modifications (top), enhancer predictions (middle) and open chromatin (bottom) in human (left) & zebrafish (right) in classes of linkage score. CNEs targeting at least one gene were divided into 10 deciles of equal size, according to their linkage scores. Class 0 represents CNEs with no associated target gene. The top row represents overlap with ChIP-seq peaks of histone modifications in human embryonic stem cells [4] or in various developmental stages in zebrafish [5]. The middle row represents overlap with enhancer predictions from FANTOM [7] or Vista [8] in human, & from differentially methylated regions during development in zebrafish [9]. The bottom row represents overlap of DNase1 peaks in human embryonic stem cells [4] and overlap of ATAC-seq regions with a normalised read count higher than 5 in zebrafish embryos [19].

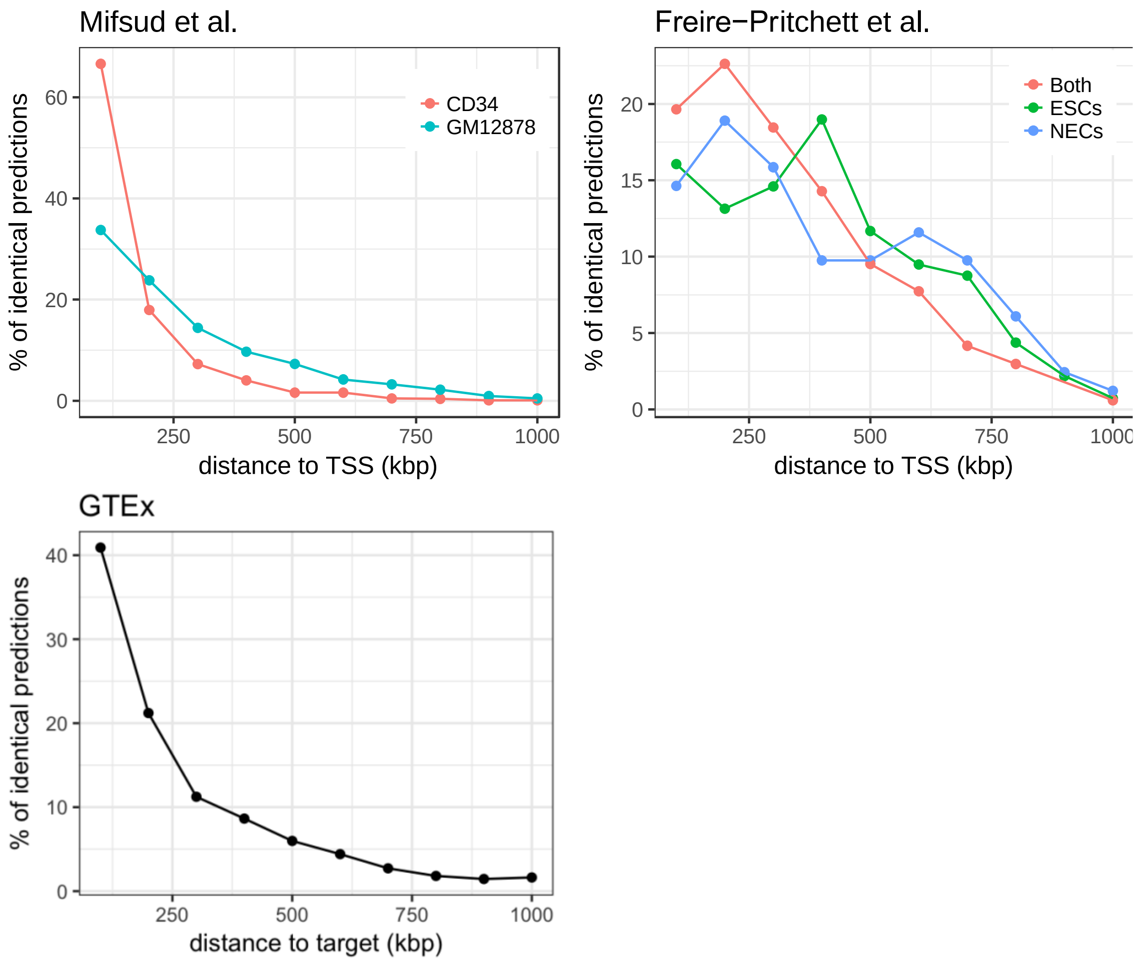


**Supplementary Figure 4:** Percentage of CNE-target gene associations in agreement with *in-vitro* experiments in classes of distances to their target's TSS. Top left & right panels: Capture Hi-C data from [11] and [10] respectively. Bottom left panel, eQTL data from the GTEx project [12]. We considered only cases where 1) a PEGASUS CNE and a predicted enhancer by another study overlapped, 2) both predicted enhancers from any method were associated with one gene only and 3) both predicted associated genes were identical.

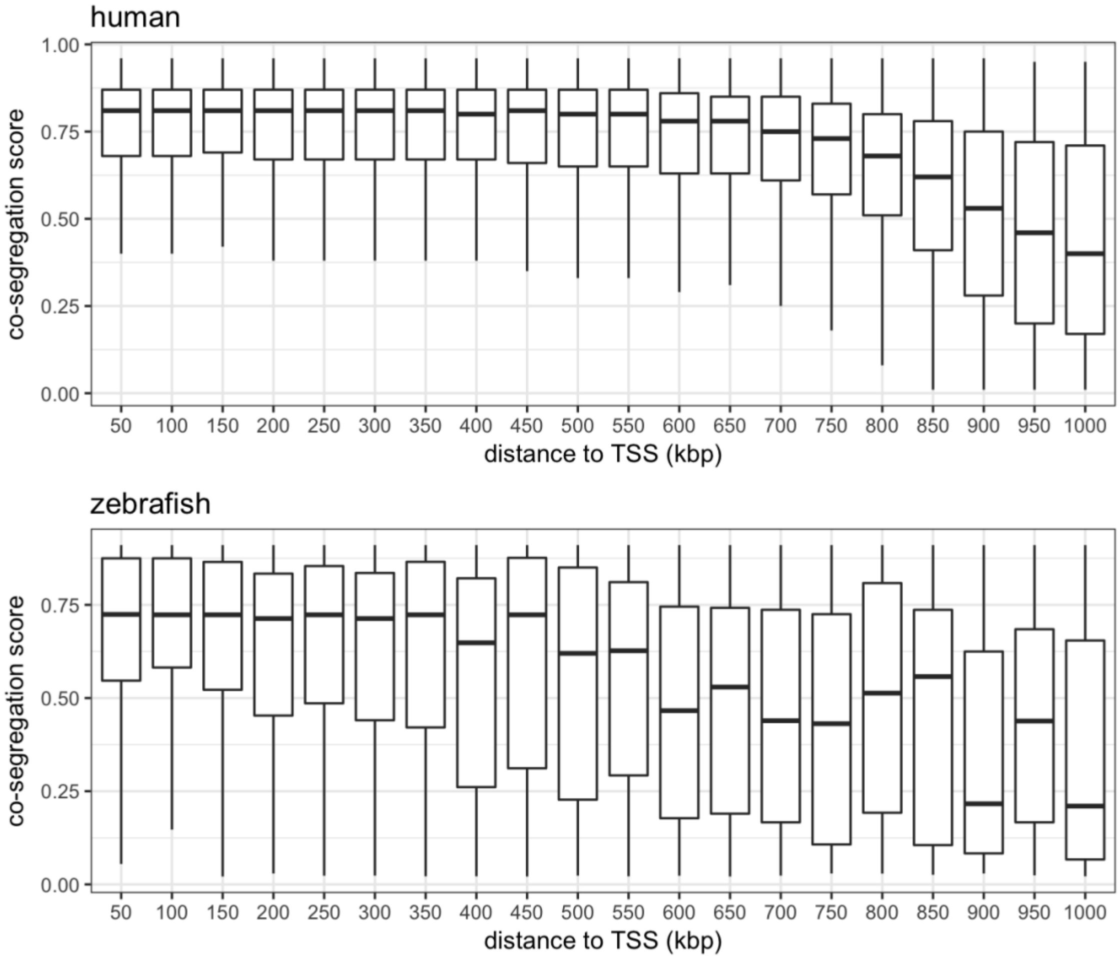


**Supplementary Figure 5:** Distributions of linkage scores as a function of the CNE-TSS distance in human (top) and zebrafish (bottom).

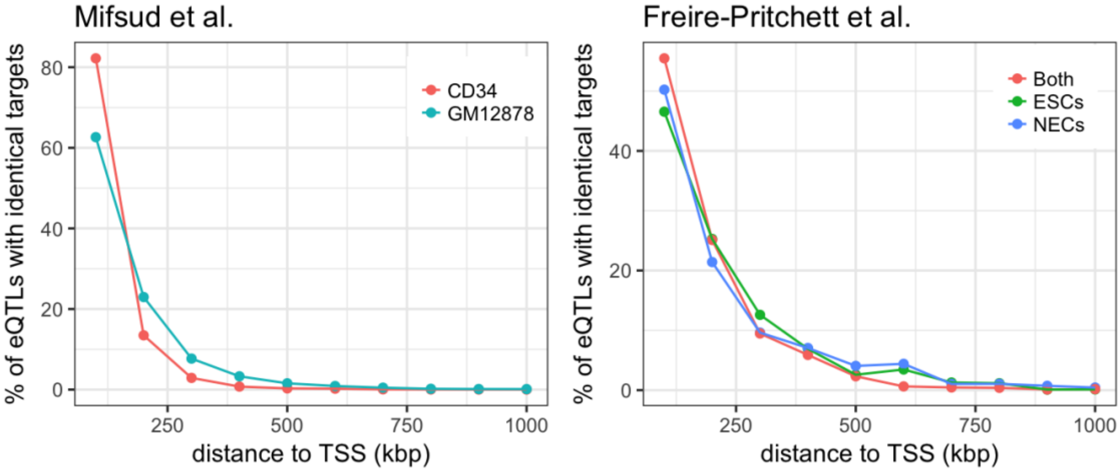


**Supplementary Figure 6.** Percentage of GTEx eQTLs target gene predictions overlapping Capture Hi-C predictions as a function of distance to target.

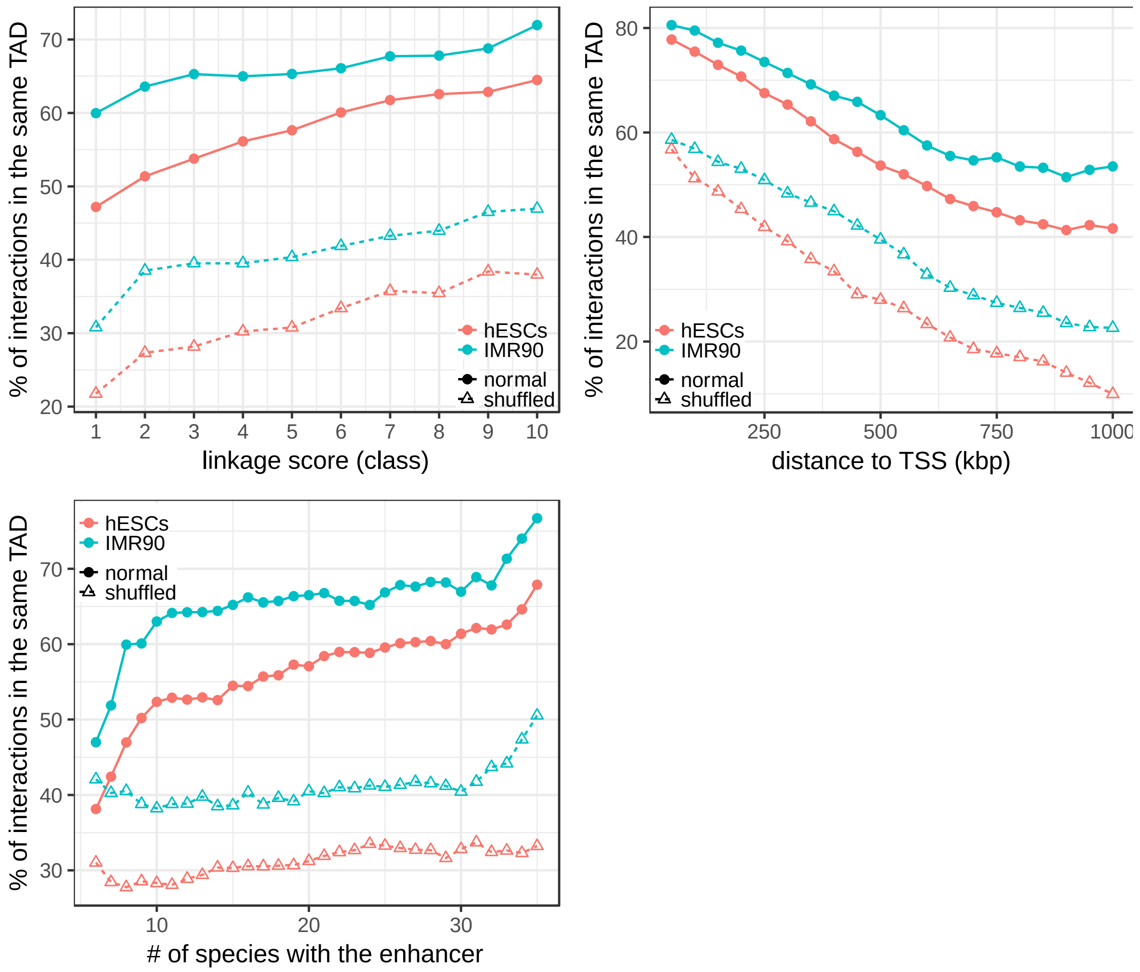


**Supplementary Figure 7:** Percentage of CNE-gene associations located in the same TAD as a function of: PEGASUS linkage score (top left panel), distance to TSS (top right panel), number of species possessing the enhancer (bottom left panel), for hESCs (red) and for IMR90 (blue) cells [13]. Dashed lines correspond to the same analysis with TADs coordinates shuffled along the human genome.

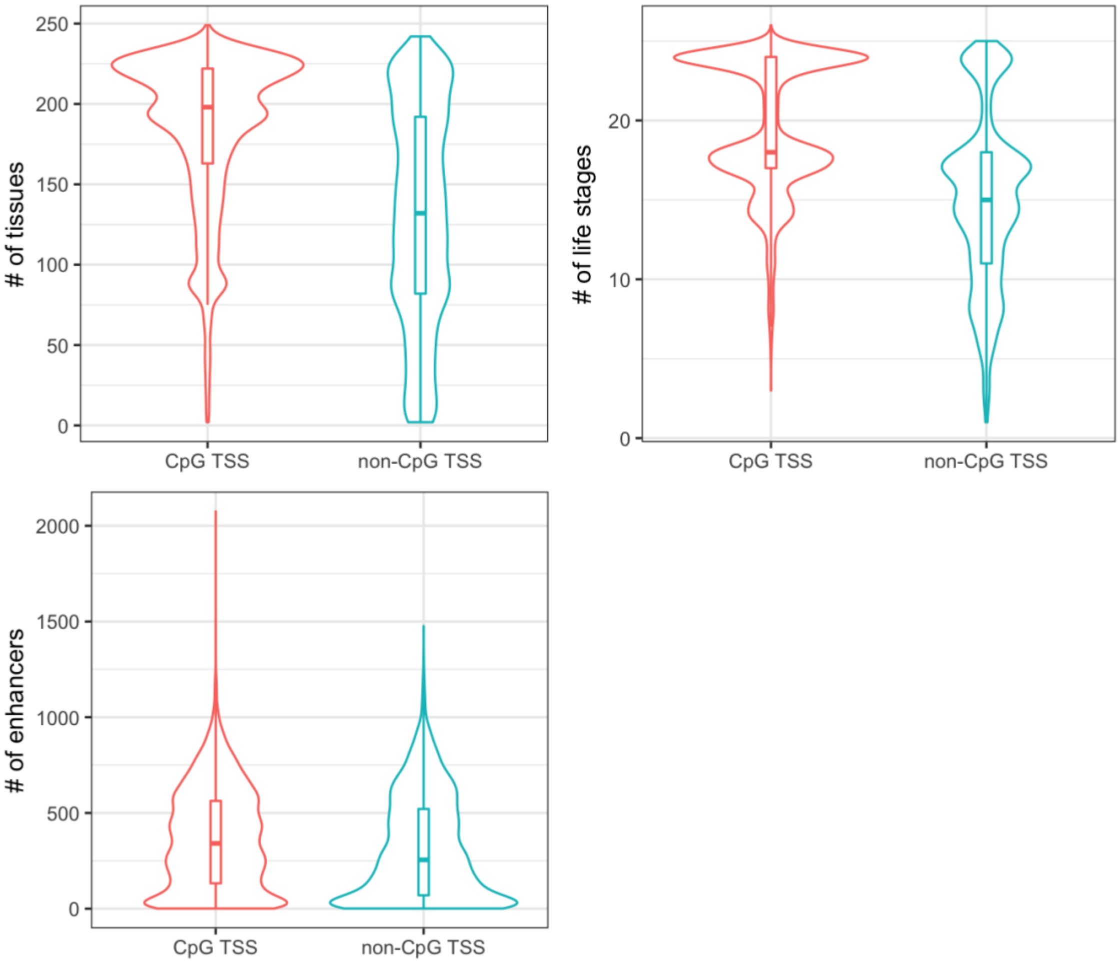


**Supplementary Figure 8:** distributions of number of tissues, life stages and CNEs for genes with TSS overlapping a CpG island (CpG TSS) and other genes (non-CpG TSS). All distribution means are different between CpG genes and non-CpG genes (Mann-Whitney-U test, all p-values < $10^{-15}$)

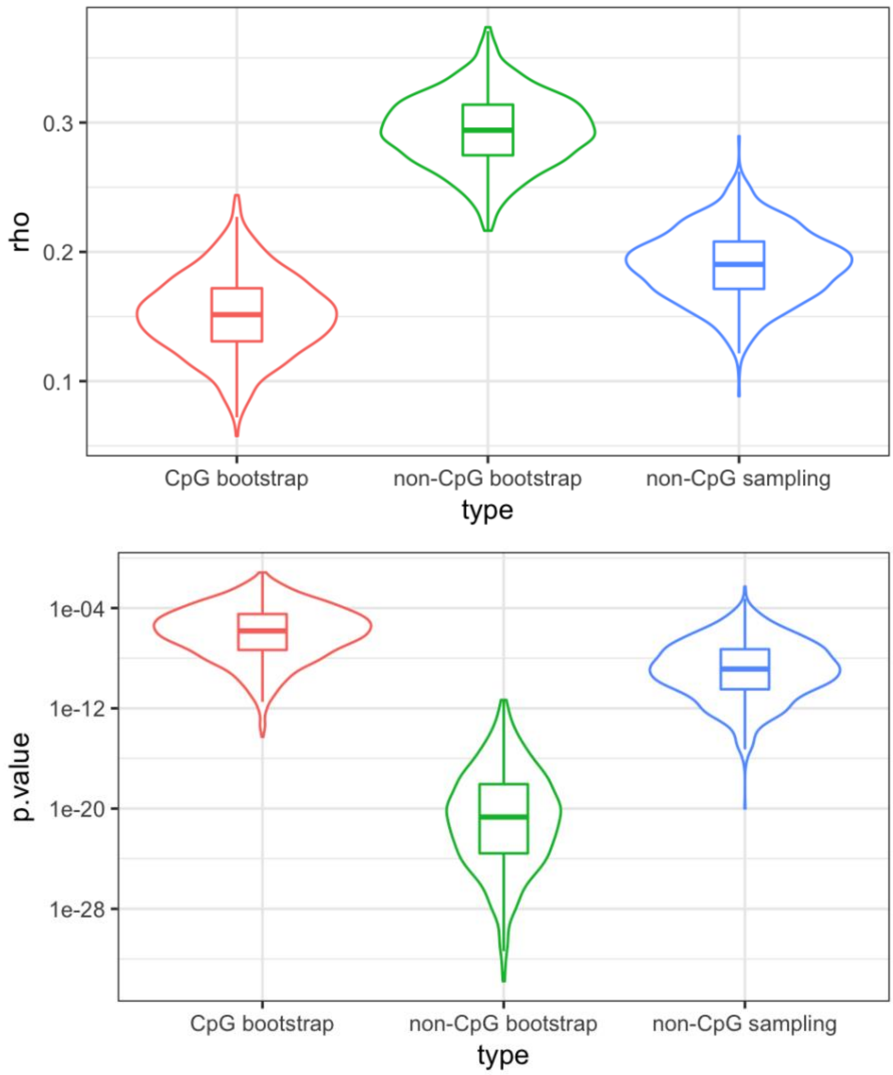


**Supplementary Figure 9:** Spearman's correlation coefficients (top) and p-values (bottom) of correlations between number of CNEs targeting a gene and number of tissues where it is expressed. CpG bootstrap & non-CpG bootstraps represent correlations in 100 bootstrap replicates for CpG genes & other genes, respectively. Non-CpG sampling represents 100 random sampling of non-CpG genes matching the distribution of tissues number of CpG genes.
